# Power Series Template Matching Model for Pitch Perception

**DOI:** 10.1101/2023.01.27.525831

**Authors:** Jun-ichi Takahashi

## Abstract

The problem of pitch perception has been the subject of a long debate between place theory and time theory. Here, we propose a Power Series Template (PoST) model to answer the problem of how and why pitch perception comes about. The sensitive measurement of acoustic signals requires efficient Sound Amplification, which inevitably accompanies mode coupling due to the perturbative non-linearities. Under reinforcement learning for Sound Localization with the second- and third-order modes as default teacher signals, a chain of coincidence is generated. After learning in Sound Localization is completed, two power series templates of 2^n^ and 3^m^ are generated, and the 2*f*_1_ − *f*_2_ coupling of the elements contained therein fills the blanks of matchable harmonics up to N=10 in the template, providing the well-known harmonic template. When complex tones containing multiple harmonics are input into the trained network, two power series are evoked from each harmonic, and from their intersection, the brain acquires the fundamental of the complex tone as the pitch. Based on this template model, consistent explanations are given for the problems of missing fundamental, pitch shift, pitch and chroma, and resolvability jump without the help of time theory.

## Introduction

Theories of pitch perception have wavered between place theory and time theory (de Cheveigné, 2004, Oxenham 2013). The theory of pitch perception began with Ohm’s resonance hypothesis and place theory (Helmholtz 1954) and was given anatomical evidence by Békésy (1949), who observed frequency-resolved auditory nerves in the cochlea. However, the fact that the fundamental in a complex tone was perceived as a pitch even when its intensity was reduced (Schouten 1940) or masked by noise (Licklider 1954) negated the simple place theory, and a theory focusing on the temporal structure was put forward. On the other hand, low-numbered, resolved harmonics were found to produce a much more robust and salient pitch than high-numbered, unresolved harmonics (Carlyon and Shackleton 1994). This produced another challenge for time theory, which typically did not predict a benefit for low-numbered harmonics over high-numbered harmonics. In the 70s, it was shown that altering the temporal structure of a sound by manipulating the phase of the harmonics that make up the complex tone had no effect on pitch perception (Wightman 1973). The idea of pitch perception as a direct physical measurement was rethought, and in the direction of considering pitch perception as pattern recognition, the pattern-transformation model was proposed by Wightman (1973), the optimum processor theory was proposed by Goldstein (1973), and a template model was proposed by Terhardt (1974). However, it was pointed out that it might be difficult to describe pitch perception by a single mechanism, because analysis by pattern matching also made it difficult to separate harmonics in higher-order complex tones, making template matching difficult (Houtsma and Smurzynski 1989). On the other hand, it was shown that fundamentals were perceived when different higher-order harmonics *nf*_0_ and (*n* − 1)*f*_0_ were presented separately to the two ears, indicating that pitch perception is not the result of physical information processing in the auditory peripheral system, but rather in the central system after the crossing of the left and right auditory nerves (Houtsma and Goldstein 1972). Revising the choice between spectral or temporal, an approach emerged that uses a flexible learning process in the neuronal system to calculate the fundamental for acoustic data that describes the two-dimensional information of frequency and temporal information as spectrograms on an equal basis. Shamma and Klein (2000) proposed a network model of pitch perception based on the coupling between harmonics due to the non-linearity response in the cochlear filter. Recently, Saddler et al. (2021) developed a deep neural network (DNN) model trained to estimate the fundamental frequency and showed that many properties of pitch perception could be quantitatively reproduced by incorporating cochlear filter properties into a DNN. However, Shamma and Klein (2000) noted the shortcomings of the template model. “The lack of convincing biological evidence so far for the existence of these templates or for how they might be generated.” The DNN model has a serious problem as it is a teleological procedure to optimize a given parameter. Sound pitch is a very limited piece of acoustic information. How much information about the object can we get from the pitch? Is the cost of creating a pitch perception system worth it? Why does the system choose the pitch as the target parameter for learning and not the center of gravity of the spectrum? Or why does the system perceive discrete harmonics without perceiving the continuous spectral envelope contrary to the Gestalt principle? The main problem with the DNN approach is that today’s DNNs are too powerful. If we consider a DNN as a transformation from an input signal to an output signal, then under the appropriate formulation of the problem and with the appropriate teacher data the DNN can give a formally accurate solution to almost any problem. The main focus of research using DNNs has shifted from how accurate transformations can be given to what internal mechanisms are responsible for high accuracy, and how they are formed.

How have organisms handled acoustic signals under survival strategies? In this paper, we consider a model of the neural network with a nonlinear input filter for pitch perception, going back to the origin of hearing. The various constraints imposed on the system cause deficiencies in the behavior in an ideal DNN and characterize the output of the DNN. We derive the system from Sound Amplification for the detection of small auditory signals and Sound Localization of Auditory Scene Analysis (ASA) (Bregman 1990). The production and operation of the system are represented by chains of discrete processes, which behave like state transitions of a finite state machine (FSM) in the mathematical modeling of computation (Hopcroft et al., 2001). We show that even under weak non-linearity limited to second- and third-order, chains of correlation learning enable the perception of higher harmonics, while at the same time multilayering of the network by chains leads to the loss of octave perception, which results in octave equivalence, and that the third-order nonlinearity, 2*f*_1_ − *f*_2_ coupling fills the deficit in the chain, which improves the pitch resolvability at lower orders and explains the resolvability jump at the 10th harmonic. We show that the template model can derive the basic properties of pitch perception without the help of time theory.

### Model

Sound signals are characterized by their amplitude and frequency. Sounds with significantly lower or higher frequencies have no invasive effect on the organism, as resonant interaction with the organism is lost. Each animal has its own species-specific perceptible frequency range. Sounds incident on the ear are converted into vibrations in the lymphatic fluid and stimulates the firing of the corresponding auditory nerves through the vibration of frequency-resolved inner hair cells. On the other hand, signals with excessive amplitude cause irreversible damage to the sensory system, so there is an upper limit to the sensitivity of hearing. Despite this, humans have a large dynamic range of up to 120 dB. It is thought that amplification in the cochlea due to active and mechanical movement in outer hair cells contributes to this (Davis 1983, Dallos 2008, Avan et al. 2012). In general, non-linear distortions often accompany the amplification with a wide dynamic range. Kadia and Wang (2003) measured the firing of neurons in open marmosets, and directly observed second- and third-order nonlinear mode coupling in the sensory response of auditory neurons in the A1 auditory cortex, showing that the central nerve uses mode-transformed harmonics and that there is strong nonlinear mode coupling that cannot be ignored (Wang 2013, Feng and Wang 2017). Aside from the sensory input, it had been known since the end of the 70s that when sound waves are incident on the ear, the ear radiates sound waves with frequencies that are two and three times the frequency of the incident sound wave. This phenomenon is called Distortion Product Otoacoustic Emission (DPOAE) (Kemp 1979, Bian and Chen 2008). The observation of second- and third-order DPOAEs is also direct evidence of non-linear mode coupling in the auditory organ and shows that the non-linear coupling of the response is sufficiently weak that it is valid to treat intermodal coupling as a perturbation. Although the DPOAE is a mechanical response of the organ to an externally emitted auditory signal and does not directly represent the sensory input, we consider it sufficiently feasible to consider that the properties of the DPOAE are quantitatively correlated with sound perception.

In response to stimulus input from the external world, animals localize objects (in which direction and how distant), judge their attribution (whether to approach, escape from or ignore), and determine their behavior. Behavior decisions are not mere reflex responses to stimuli, but are considered to be the output of a simulation in a model of the external world (representation) according to the location and attribution of the object in the brain through segregation and integration to the input signal. The process of reconstructing the placement of an object in the brain from acoustic signals is called Auditory Scene Analysis (ASA). The localization of objects by sound signals is called Sound Localization. As sounds of various frequencies arrive at the ear at different times, Sound Localization is the process of performing ASA by segregating and integrating sounds emanated by objects, using sound intensity, binaural time differences, and other factors as keys.

Based on these previous studies, we propose a model for creating the auditory network through the following processes. In the undifferentiated early stages of hearing, auditory cells are randomly distributed. Signals from multiple auditory cells are sent to neurons to measure their coincidence which is used to segregate and integrate sounds. Learning in neurons is considered to follow the Hebbian rule. That is, when signal input is superimposed in the presence of teacher signals, the neuron undergoes reinforcement learning. The second- and third-order overtones described above always satisfy the coincidence with the fundamental, so they act as pre-prepared teacher signals in learning. Furthermore, when a sound matching these overtones is superimposed from the outside, the neuron undergoes reinforcement learning for the correlations between the fundamental and these overtones. The reinforcement learning is bidirectional, so for each frequency *f*, correlations are implemented not only with 2*f* and 3*f*, but also with 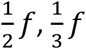 and their combinations 2^*n*^3^*m*^*f* over any integer order (-∞<n, m<∞). (Fig.1)

**Figure 1.**
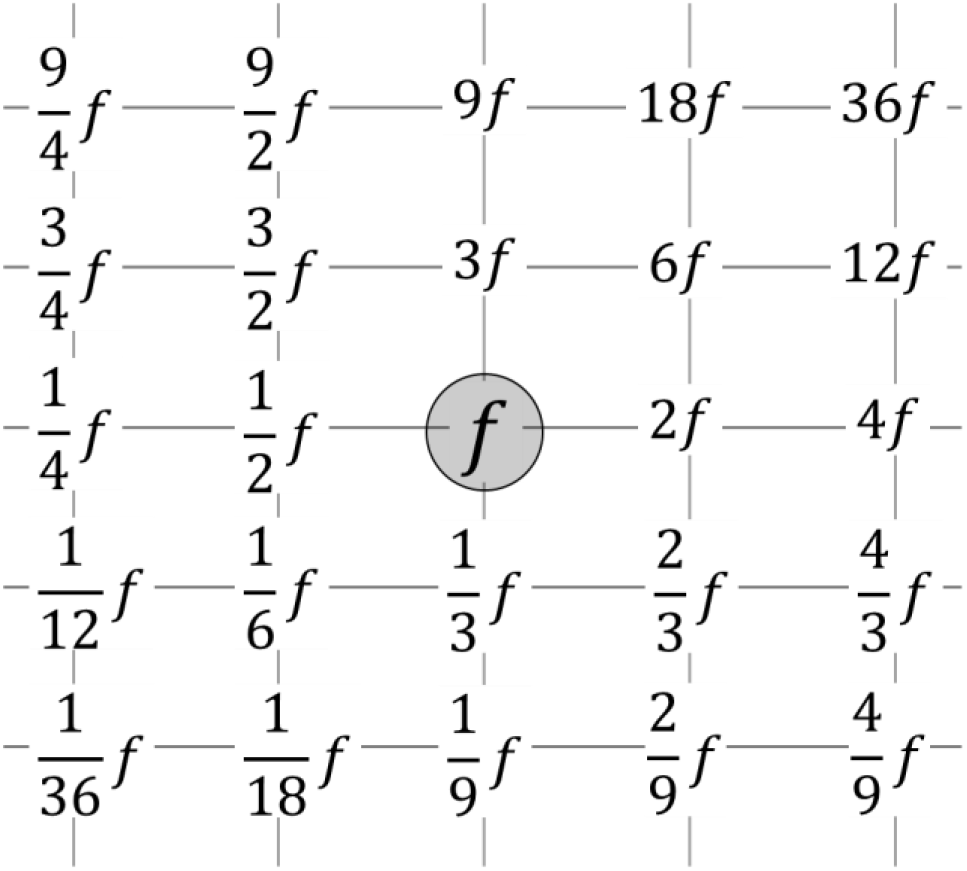
Network of evoked signals. Note: When a sound with frequency *f* is heard, neurons associated with fire simultaneously. Under the third-order nonlinearity of 2*f*_1_ − *f*_2_, 5*f* and 7*f* neurons can be fired by the recursive interaction from 2^*n*^3^*m*^*f* neurons. If a sound with 2^*n*^3^*m*^*f* is overlapped, the coincidence with *f* is perceived.

Since learning is performed in parallel for all frequencies {*f*^(*k*)^} consisting of the input sound, where *f*^(*k*)^ = *kf*_0_ with the fundamental frequency *f*_0_ and harmonic number *k*=1 to ∞, power series matching templates, [⟨2^*n*^3^*m*^*f*⟩] for every frequency *f* are formed on the distributed neurons. The neurons in the series fire simultaneously for all individual inputs, so frequency discrimination within it is not possible. At the same time, as described below, the third-order nonlinearity 2*f*_1_ − *f*_2_ adds 5th and 7th matching template harmonics. Finally, harmonic templates, [⟨2^*n*^3^*m*^*f*⟩, ⟨5*f*⟩, ⟨7*f*⟩] are produced. The first column in Table 1 shows the harmonics linked by chains of second- and third-order nonlinearities. Here n=5,7 are prime to 2,3 and each other, so they cannot be directly linked by second- and third-order harmonics with the fundamental. On the other hand, under third-order nonlinearity, a third wave 2*f*_1_ − *f*_2_ is generated from two different waves of *f*_1_ and *f*_2_. Consider this three-wave pair: when two sounds with *f*_1_ and *f*_2_ are input, a signal with a frequency of 2*f*_1_ − *f*_2_ acts as a prepared teacher signal. When a sound with 2*f*_1_ − *f*_2_ are superimposed simultaneously, the neuron undergoes reinforcement learning of the coincidence among the three waves. After learning, when any two of *f*_1_, *f*_2_ and 2*f*_1_ − *f*_2_ are input, regardless of the order of frequency, signals with the remaining frequency is perceived simultaneously. The blank n=5, 7 in the column of Table 1 can be learned to correlate with the harmonics in the second and third columns by 5=2*4-3, 7=2*8-9, respectively. Similarly, n=10, 14, and 15 can be learned by the coincidences with the harmonics in the second and third columns by 10=2*9-8, 14=2*16-18, and 15=2*12-9, respectively. Although n=14, 15 can also be decomposed into 2*7, 3*5, the learning efficiency would be lower because 2*f*_1_ − *f*_2_ learning requires two teacher signals whereas overtone learning requires only one externally provided teacher signal, and the joint strength of the coincidences learned in 2*f*_1_ − *f*_2_ learning is considered weaker than in overtone learning. Therefore, for n=14 and 15, matching 14=2*16-18 and 15=2*12-9 are considered to be preferred. Also, n=11, 13, 17, and 19 are prime to 2 and 3 each other and cannot be factorized by 2 and 3. It seems that 2*f*_1_ − *f*_2_ learning is possible using the second- and third-column harmonics as these are decomposed, e.g., 11=2*6-1, 13=2*8-3, 17=2*9-1. However, DPOAE shows a sharp decrease in coupling strength for *f*_1_/*f*_2_>1.4 (Bian and Chen 2008). Considering the contribution of amplification by outer hair cells, learning at *f*_1_/*f*_2_>1.4 seems to be difficult. Therefore, the harmonics learned by the system should be restricted to the numbers without brackets in the first column of Table 1. Once the learning is established, the system acts as a coincidence detector between the fundamental and the harmonics. The discrete internal structure acquired after the convergence of learning with statistical data will give the system robustness to minor variations in details of the linear properties of input filters, fluctuations in network parameters, etc. We name the harmonic template system derived from the chain of correlation learning Power Series Template (PoST) system. Here, the PoST system includes the 5th and 7th harmonics through the 2*f*_1_ − *f*_2_ coupling. As mentioned above, learning the 2*f*_1_ − *f*_2_ coupling is less efficient than direct power series learning. If it is necessary to consider a PoST system without 5th and 7th harmonics, we call the system pre PoST (Fig.1). In the inference process of the PoST system, the input acoustic signal undergoes a step-by-step transformation to a pitch, as will be shown in the next chapter. The production and operation of the PoST system are depicted in Fig.2. The step-by-step transformation can be considered an operation in an FSM if it is regarded as a transition between states. The system acts as a discrete signal processor and the FSM provides a simple description of operations that is robust for small fluctuations in the parameters. In the following, the operation of the PoST system and its characteristics are described in detail.

**Figure 2.**
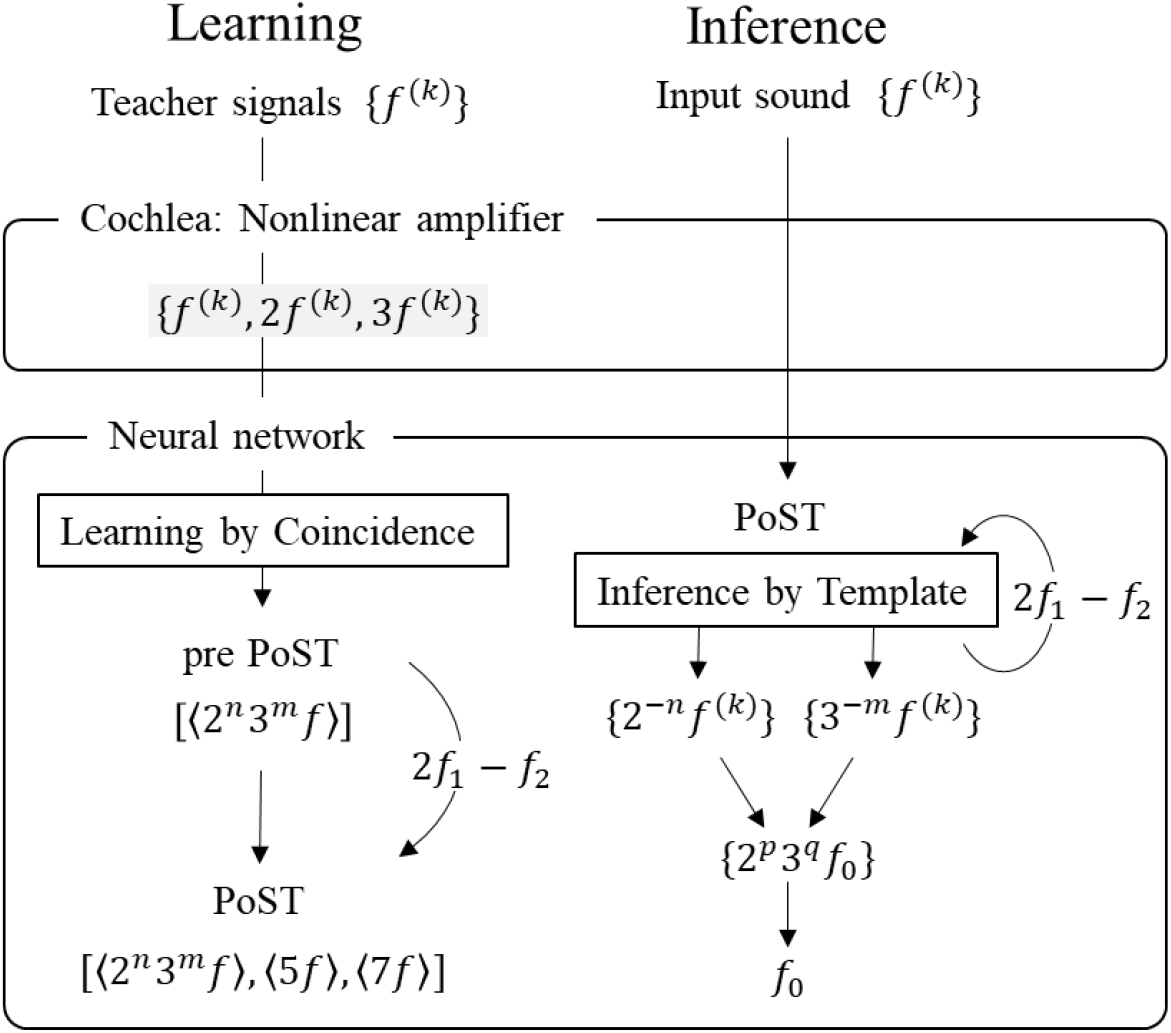
Learning and inference of PoST system. Note. {*f*^(*k*)^} are harmonic series of *kf*_0_ with a fundamental frequency *f*_0_ and k=1 to ∞. {*f*^(*k*)^, 2*f*^(*k*)^, 3*f*^(*k*)^} are combined series with second- and third-order harmonics. [⟨2^*n*^3^*m*^*f*⟩] are power series harmonic templates for each *f*. [⟨2^*n*^3^*m*^*f*⟩, ⟨5*f*⟩, ⟨7*f*⟩] are the PoST template to which 5th and 7th matching harmonics are added. In the learning process, nonlinear cochlear amplification provides second and third harmonics and they are exploited by the NN as teacher signals, producing PoST. In the inference process, PoST outputs chains of the power series of second- and third-order couplings from each *f*^(*k*)^. If they have an overlap, they are integrated into a single series with the fundamental frequency *f*_0_. If *f*_0_ is consistent with all of *f*^(*k*)^, *f*_0_ is output as the fundamental frequency of the input sound.

**Table 1.** Harmonic template learned by the second- and third-order nonlinearities.

| N | $2^n$ | $3^m$ | $2f_1 - f_2$ | N | $2^n$ | $3^m$ | $2f_1 - f_2$ |
| --- | --- | --- | --- | --- | --- | --- | --- |
| 1 | | | | 14 | | | $2*16-18$ |
| 2 | 1 | | | 15 | | | $2*12-9$ |
| 3 |  | 1 |  | 16 | 4 |  |  |
| 4 | 2 |  |  | (17) |  |  |  |
| 5 | | | $2*4-3$ | 18 | 1 | 2 | |
| 6 | 1 | 1 |  | (19) |  |  |  |
| 7 | | | $2*8-9$ | 20 | | | $2*18-16, 2*16-12$ |
| 8 | 3 |  |  | (21) |  |  |  |
| 9 |  | 2 |  | (22) |  |  |  |
| 10 | | | $2*9-8, 2*8-6$ | (23) | | | |
| (11) |  |  |  | 24 | 3 | 1 |  |
| 12 | 2 | 1 |  | ... |  |  |  |
| (13) |  |  |  |  |  |  |  |

Note. First column: the order of harmonics, second and third columns: the number of iterations of second-order and third-order nonlinear coupling required to access the harmonics, fourth column: the combinations of those defined from the second and third columns to access the harmonics by the $2f_1 - f_2$ coupling. Here, those of $f_1/f_2 > 1.4$ are not written (see text). The numbers in brackets are not accessible by the second- and third-order nonlinearities.

## Discussion

### Missing fundamental

It has been a fundamental problem in pitch perception that fundamental missing harmonic sequences induce pitch sensation (Seebeck 1841, Schouten 1940, Licklider 1954).

Examples of the matching scheme of the PoST system is shown in Fig.3. When a sound consisting of multiple harmonics {*f*^(*k*)^} is input to a trained neural system, 2^n^ and 3^m^ series are evoked separately for each harmonic and are cross-checked for coincidence. (Here, only two tones are shown in Fig.3.) If there is an intersection between the two series, the frequency of the intersection is the fundamental for the input harmonics (Fig.3a). Even when there is no intersection between the respective series of the two input harmonics, the third-order nonlinearity evokes 2*f*_1_ − *f*_2_ and 2*f*_2_ − *f*_1_ harmonics from *f*_1_and *f*_2_, which gives another path for matching in the PoST (Fig.3b). If their networks have an intersection *f*_0_ with the network of *f*_1_ or *f*_2_, the neuronal system returns *f*_0_ as the missing fundamental and *f*_1_ and *f*_2_ are written as the harmonics of *f*_0_. When the input sound has more than three harmonics, matching is performed in parallel in all pairs of auditory neurons. However, it is not necessary to complete matching on every pair. If a harmonic template generated by the fundamental determined from any two harmonic pairs is consistent with the rest of the harmonic group, no further matching is required, and the Sound Localization can be terminated. The determination of the pitch signal would give a cue to take the exit action while matching is not completed in other neurons, which would act on the auditory system to interrupt attention to the object and prevent it from wasting further resources for perception. It is noted that the fundamental is not prepared a priori as a generator of the harmonic template (and is sometimes missing in the auditory input), but rather it is more appropriate to think of it as being defined recursively from the input harmonics. The system does not perceive the auditory signal as a whole spectrum, but integrates it into a special single scalar quantity, the pitch of the sound, which allows the target signal to be incorporated into the ASA with minimal resources in perception just as the edge detection in vision. The parsimony would be critical for quick behavior decisions.

**Figure 3.**
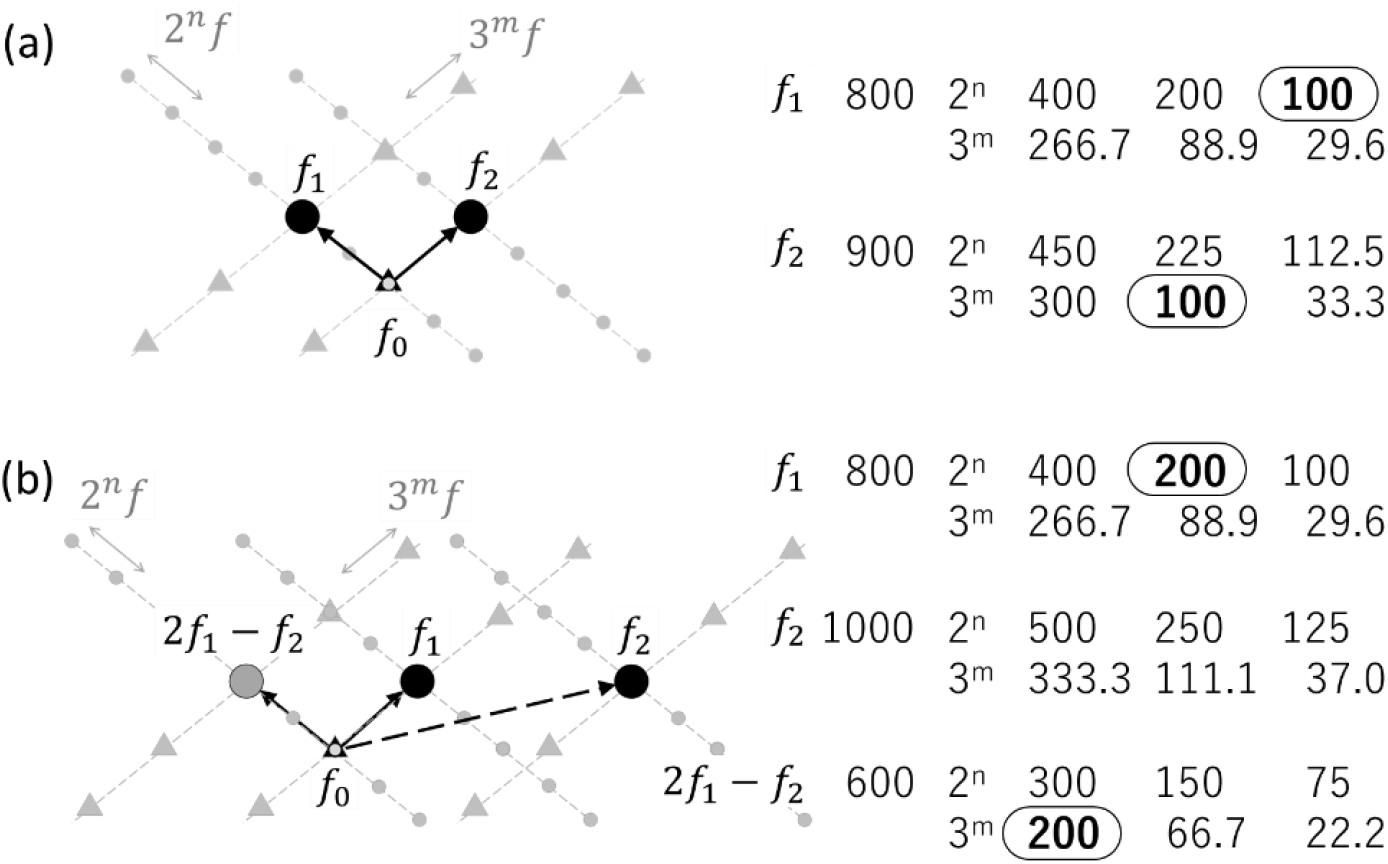
Matching scheme of PoST system. Note. (a): when 800 Hz and 900 Hz tones are input simultaneously, each evokes 2^n^ and 3^m^ power series templates, respectively. The intersection 100 Hz (bold) is uniquely determined from the former 2^n^ template and the latter 3^m^ template. (b): when the 800 Hz and 1000 Hz tones are input simultaneously, there is no intersection in the power templates evoked from each. However, they evoke a 600 Hz tone by the 2*f*_1_ − *f*_2_ coupling, defining the intersection 200 Hz (bold) uniquely from the 2^n^ template of 800 Hz and the 3^m^ template of 600 Hz.

### Pitch shift

When all of the components in a harmonic complex tone are shifted in frequency by Δ*f*, the perceived pitch of the complex shifts roughly in proportion to Δ*f*. The proportionality factor can be approximated by the reciprocal of the harmonic order of the carrier frequency of the complex tone regarding the pitch fundamental frequency (Schouten 1940, Schouten et al. 1962).

The learning of the power series template is carried out in parallel at all frequencies. When a frequency is fixed, the tuning width of its harmonics learned by power harmonics cannot be infinitesimally small and always has a finite width. Once the learning is complete, matching at the higher harmonics with a detuning is possible. In listening to complex tones composed of multiple harmonics of a common fundamental, when the harmonics are shifted by the same frequency, the fundamentals determined by power series template matching for each harmonic will have different values. However, if the modulation is within the tuning width, the brain will prefer to match them with a common frequency. In a trained network, the firing probability of a neuron continuously decreases with greater detuning. Approximating the likelihood for detuning by the simplest additive quadratic function, for example, the likelihood in a three-tone complex is 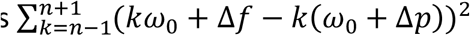, and the best estimate is 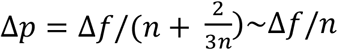 (the first effect of pitch shift), where *kω*_0_ + Δ*f* is the *k*th harmonic including the frequency shift Δ*f* used in the experiment, and *ω*_0_ + Δ*p* is the virtual fundamental. It is known that the proportionality factor obtained experimentally deviates systematically from 1/*n* (the second effect of pitch shift) (Schouten et al. 1962, Smoorenberg 1970). Its origin could be attributed to the characteristics of the likelihood function.

### Pitch and chroma

It is well known that sounds with twice the frequency are perceived as having the same pitch. (Octave circularity in pitch perception) (Deutsch 2010, Shepard 1982).

In our model, harmonics that differ by a factor of 2^n^ in frequency are integrated into a single series during learning and are no longer distinguishable. This is consistent with octave circularity. Similarly, harmonics with different 3^m^ fold frequencies are integrated into a single series, which is known as the perfect 5th consonance. Shepard has geometrically represented the periodicity including octave and perfect 5th in pitch with the double helix model (Shepard 1982). In general, the harmonics of the sounds that make up the object sound consist of harmonics of the natural number of orders of the fundamental harmonic. On the other hand, the power series templates contain harmonics with negative integer powers, and the perception of subharmonics would be learned simultaneously due to the bidirectionality of the learning, whereas they do not contribute to the Sound Localization. The subharmonics can be folded back to a region above the fundamental frequency by multiplying by 2^n^ or 3^m^. These are frequencies with a rational ratio to the fundamental and define a chroma. Which chroma is preferred or survives will depend on culture, and resulting musical scales.

### Resolvability

The fundamental discrimination threshold (F0DT) of the complex harmonics tone, which consists of several harmonics, shows a critical threshold around N=10 (Houtsma and Smurzynksi 1990, Bernstein and Oxemham 2003). Bernstein and Oxemham investigated it using amplitude-modulated tones consisting of three consecutive harmonics, showing high discrimination performance at N≦10, with a sharp drop at N>10. However, there is no significant change in discrimination performance at much higher orders. Because the components of a harmonic complex are equally spaced on a linear frequency scale, but the absolute bandwidths of auditory filters increase with increasing center frequency, the density of harmonics per auditory filter increases with increasing harmonic number. As a result, low-order harmonics are resolved from one another, but higher-order harmonics begin to interact within single auditory filters and eventually become unresolved. Then, it was proposed that the pitch perception was attributed to matching with the harmonic template for low-order harmonics and reading the period pattern of the acoustic waveform for higher-order harmonics.

This discontinuous change has been one of the biggest problems in pitch perception theory. Let us look at this problem from the standpoint of power series template matching. Consider a complex tone consisting of several harmonics (Table 2). If the tone contains two consecutive orders of harmonics, the 2*f*_1_ − *f*_2_ coupling produces the perception of two harmonics below and above the orders at the same time (underlined). A power series template matching is performed among a total of four harmonics. In each complex tone, the 2^n^ and 3^m^ power series template intersections are calculated consistently by the harmonics in bold, which give matching solutions listed in the third column. The matching calculation has a solution for all combinations up to N≦10. In this case, it is not necessarily the fundamental that is determined by the calculation, but the doubling for the pairs (5,6) and (6,7) and the tripling for the pairs (10,11). The multiples do not match the original harmonic pairs. They are folded again into the fundamental to complete the template matching.

**Table 2.**
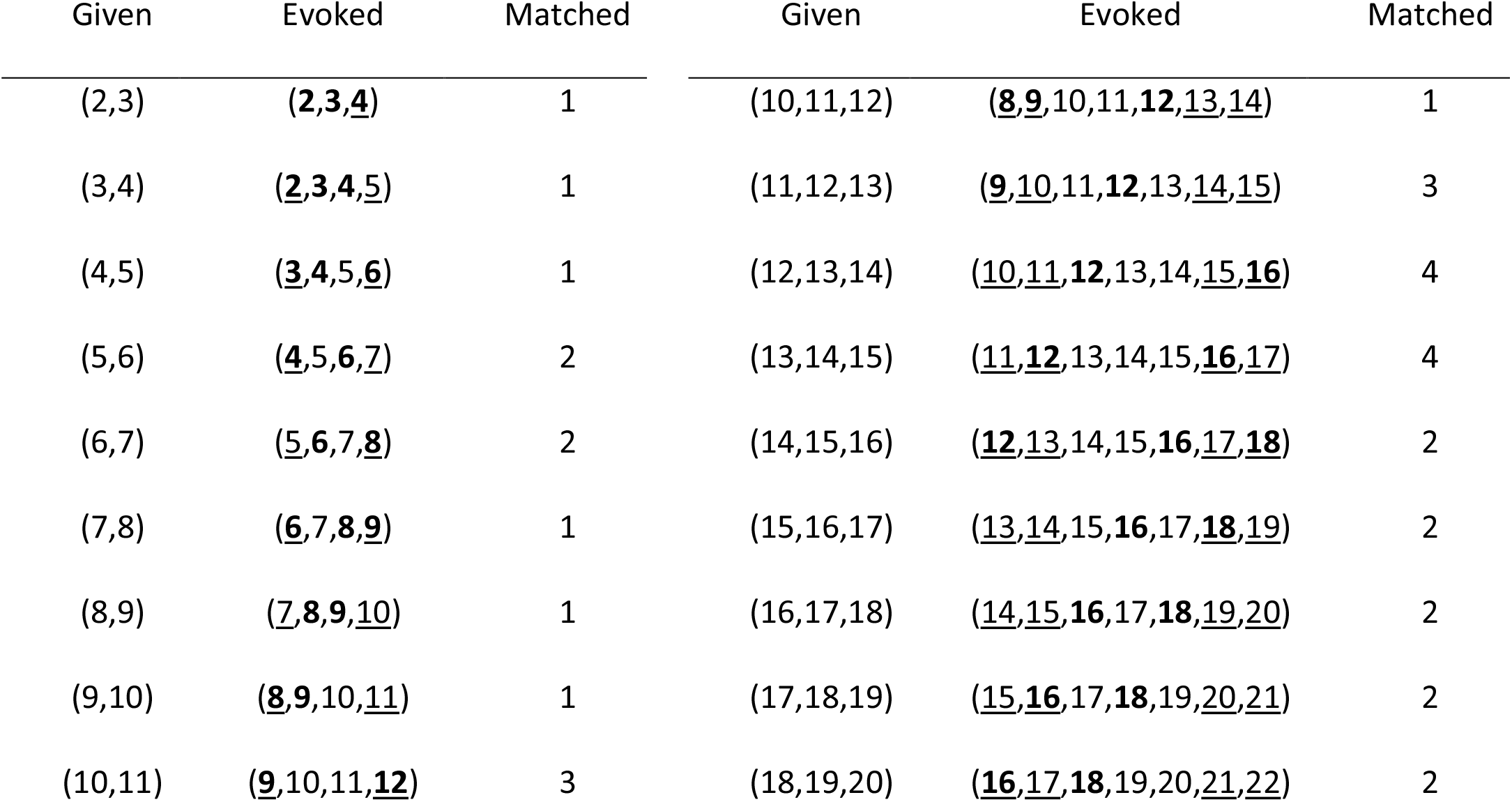
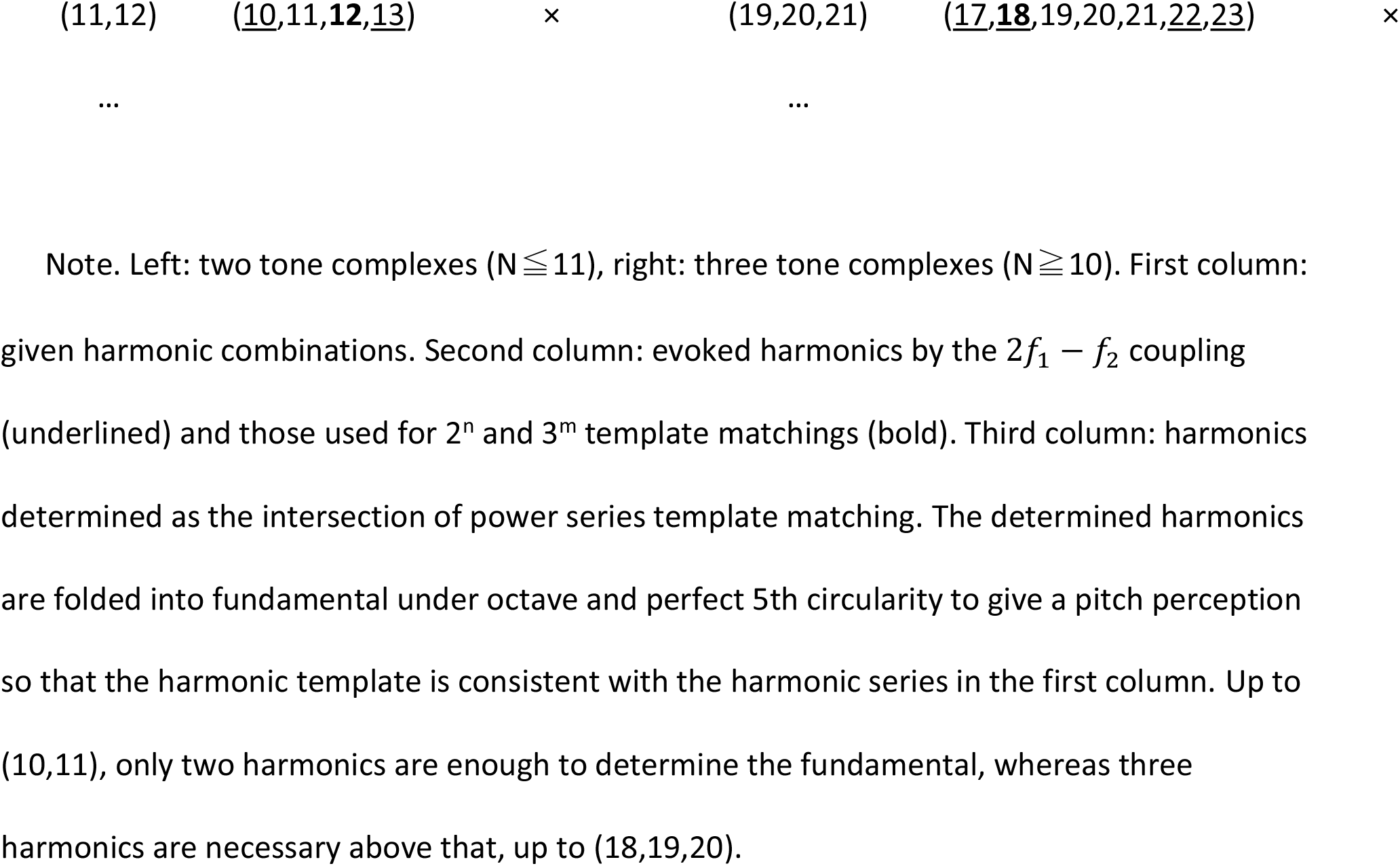
Calculation of fundamentals in complex tones with consecutive harmonics.

Even if the complex tone contains more harmonics, if they match a harmonic template, they will be integrated into the same power series templates, giving the same fundamentals. The matching calculation using two harmonics except (16,17) and (17,18) does not have a solution for N≧11, but by using three consecutive harmonics, it is possible to have solutions up to (18,19,20). An increase in the number of harmonics for matching calculation decreases the efficiency of the matching calculation and therefore the fundamental discrimination thresholds for N≧11 would have to be larger than that for N≦10. Our answer to the problem of resolved and unresolved harmonics for pitch perception is the special structure of the matching template itself rather than the transition to the perception of time-structure.

## Conclusion

We have approached the problem of pitch perception with the help of nonlinear dynamics of the cochlear amplifier, evolution of hearing, auditory psychology based on ASA, and discrete mathematics in it. We started from perturbative second- and third-order nonlinearity of cochlear amplification of small auditory signals, and considered the chains of the reinforcement learning of neural network, which produced a power series harmonic template. The third-order nonlinearity, 2*f*_1_ − *f*_2_ coupling filled some blanks in the raw-power series harmonic template, which gave the harmonic template system named PoST. The chains enabled the perception of coincidence between the fundamental and higher harmonics at the cost of losing the octave perception, resulting in the octave equivalence. The 2*f*_1_ − *f*_2_ coupling filled the 5th and 7th blanks of the raw-power series harmonic template, resulting in an improvement of F0DT for complex tones below 10th. We attributed the characteristic features of pitch perception to the constraints on the ideal DNN. Our model could give not only explanations about the characteristic features of pitch perception without the help of time theory but also the answers to fundamental problems of pitch perception. What is pitch perception? The success of Sound Integration of emanated sound from a single object. Why was pitch perception born? Reinforcement learning of Sound Localization using second- and third-order nonlinearity as teacher signals. Why is the pitch classified into discrete chroma? Subharmonic perception from the bidirectional coupling in harmonic folding.

### Open practices statement

Neither of the studies reported in this article has been pre-registered. No data other than those presented here are available.

